# APRIL drives a co-ordinated but diverse response as a foundation for plasma cell longevity

**DOI:** 10.1101/2021.06.15.448496

**Authors:** Sophie Stephenson, Matthew A Care, Gina M Doody, Reuben M Tooze

**Affiliations:** Division of Haematology and Immunology, Leeds Institute of Medical Research, University of Leeds, Leeds, LS9 7TF, UK; Bioinformatics Group, School of Molecular and Cellular Biology, University of Leeds

**Author notes:** These authors contributed equally to this work (Sophie Stephenson, Matthew A Care). Corresponding author: Reuben Tooze.

**Keywords:** Human, B cells, Plasma Cell, Plasmablast, Antibodies, Cell Activation, APRIL, MYC, Cell growth, Myeloma

## Abstract

Antibody secreting cells (ASCs) survive in niche microenvironments, but cellular responses driven by particular niche signals are incompletely defined. The TNF superfamily member APRIL provides a niche signal that can support the transition of transitory plasmablasts into long-lived plasma cells. Here we explore how APRIL helps to establish the biological programs that promote life in the niche, by studying the initial response of primary human plasmablast to APRIL. Under conditions allowing the maturation of ex vivo or in vitro generated plasmablasts, we find that APRIL drives activation of ERK, p38 and JNK. This is accompanied by a classical NFκB response. Under these conditions induction of AKT phosphorylation is also observed with similar kinetics, paralleled by FOXO1 phosphorylation and nuclear exclusion. Time course gene expression data resolve the downstream co-ordinated transcriptional response. The APRIL-signal propagates via immediate early genes and classical NFκB responsive targets to converge onto modules of MYC- and OCT2-regulated gene expression linked to cell growth, as well as leading to enhanced expression of ICAM1 and SQSTM1 associated with adhesion and metabolic/stress responses. Thus, APRIL drives a combination of multiple transcriptional programs that co-ordinate cell growth, stress response and adhesion in human ASCs, providing a broad foundation to support plasma cell longevity.

## Introduction

The survival of plasma cells (PCs) is dependent on specific niche conditions. On the one hand this allows the maintenance of long-lived humoral immunity, and on the other hand provides a flexible mechanism for limiting the PC pool.(1) Additionally, both the nature of the differentiating B-cell, the type of signal driving differentiation and the nature of the niche in which the antibody secreting cell (ASC) eventually survives as a long-lived PC (also referred to as memory PCs) may convey functional specialization.(2, 3) Several niche factors have been defined which may contribute to the survival of an ASC and allow the maturation from the transitional plasmablast (PB) state, which couples proliferation and acquisition of secretory capacity, to the quiescent but long-lived PC state.(1, 4–6) However, relatively little is known regarding the usage of specific signaling pathways and the downstream transcriptional responses to specific niche signals in ASCs. Here we use a model system which allows the in vitro generation of long-lived human PCs to study the response of human ASCs to the niche factor APRIL, as the cells initiate the final differentiation step to the quiescent PC state.

APRIL belongs to the TNF-superfamily (TNFSF). This superfamily along with its cognate receptors, includes several critical regulators of B-cell survival, activation and commitment to the ASC differentiation fate.(7) APRIL and its most closely related TNFSF member BAFF share partially overlapping receptors in BCMA (TNFRSF17), TACI (TNFRSF13B) and BAFFR (TNFRSF13C).(8) These receptors are themselves regulated during differentiation of B-cells, such that BAFFR dominates in resting B-cells allowing effective signaling from BAFF but not APRIL, while BCMA predominates in PBs and PCs. TACI bridges these patterns with expression peaking during activation.(7) This provides the potential for preferential responses from APRIL rather than BAFF at later stages of differentiation. Further layers of regulation operate both in relation to the shedding of surface receptors, and the extent of oligomerization of the ligands.(9) Notably BCMA, the primary receptor for APRIL, can be cleaved and shed from the cell surface by the action of γ-secretase, which has recently been identified as a limiting factor for APRIL responses in PC populations in vivo and in cell lines in vitro.(10)

The importance of APRIL/BCMA signals to PC survival has been indicated both in murine knockouts(11, 12) and in humans where targeting has been explored as a therapeutic avenue in rheumatological conditions.(13, 14) In both contexts the data support the conclusion that signals delivered by these factors are required for optimal PC survival. Functionally, in murine PCs BCMA signals supports survival through induction of MCL1(15) and BCMA signals can also support myeloma cell survival in vitro and in vivo.(16) Studies of signaling have demonstrated the activation of MAP kinase pathways and classical NFκB responses in B-cells following stimulation with BAFF and in cell line models of PC neoplasia in response to APRIL.(7, 17) In heterologous expression systems BCMA signaling has been shown to have the potential to activate p38, JNK MAP kinase pathways alongside classical NFκB responses.(18) Indeed, in PC neoplasia mutations affecting the NFκB pathway, are frequent events associated with progression and independence from niche survival signals, potentially substituting in part for APRIL/BCMA signals.(19–21) While APRIL has been shown to provide a survival signal for PCs in vitro and in vivo, the signaling and immediate gene regulatory responses triggered by this factor in human ASCs have not to our knowledge been explored in detail.

Here we have addressed the question of how human ASCs respond to the APRIL niche signal focusing on the responses that occur at the transition between proliferating PB and quiescent PC. Under conditions that efficiently promote the survival of both in vitro generated and ex vivo derived human PBs, APRIL drives a selective pattern of MAP kinase activation alongside classical NFκB signaling and AKT activation. The APRIL response induces a series of gene expression changes including waves of immediate early and secondary response genes, which converge onto MYC and cell growth modules of gene expression and also include changes in genes such as *ICAM1* related to cell adhesion. Thus, while promoting survival, APRIL delivers a complex signal potentially promoting growth as well as adhesion in PCs.

## Materials & Methods

### Reagents

IL2, IL21, IL6, IFNα (Miltenyi); Multimeric-APRIL H98, Multimeric-CD40L (AdipoGen); Goat anti-human IgM & IgG F(ab’)_2_ fragments (Jackson ImmunoResearch); Lipid Mixture 1, chemically defined (200X) and MEM Amino Acids Solution (50X) (Sigma); L-685,458 (gamma-secretase inhibitor, GSI) (Tocris).

### Donors and cell isolation

Peripheral blood was obtained from healthy donors after informed consent. The number of donors per experiment is indicated in the figure legend, with each symbol representing a different donor. Mononuclear cells were isolated by Lymphoprep (Abbott) density gradient centrifugation. Total B-cells were isolated by negative selection with the Memory B-cell Isolation Kit (Miltenyi).

Peripheral blood samples from anonymous donors were obtained on day 6-8 post influenza vaccination (2017-18). BCMA positive cells were isolated using a combination of BCMA-biotinylated antibody and anti-biotin microbeads (Miltenyi).

### Cell cultures

Cells were maintained in Iscove’s Modified Dulbecco Medium (IMDM) supplemented with Glutamax and 10% heat-inactivated fetal bovine serum (HIFBS, Invitrogen); Lipid Mixture 1, chemically defined and MEM Amino Acids solution (both at 1x final concentration) were added from day 3 onwards.

Day 0 to day 3 – B cells were cultured in 24 well plates at 2.5 × 10^5^/ml with IL2 (20 U/ml), IL2l (50 ng/ml), F(ab’)_2_ goat anti-human IgM & IgG (2 μg/ml) on γ-irradiated CD40L expressing L-cells (6.25 × 10^4^/well).

Day 3 to day 6 – At day 3, cells were detached from the CD40L L-cell layer and reseeded at 1 × 10^5^/ml in media supplemented with IL2 (20 U/ml) and IL21 (50 ng/ml).

Day 6 to day 13 – At day 6, cells were harvested and seeded at 1 × 10^6^/ml in media supplemented with IL6 (10ng/ml), IL21 (10 ng/ml), GSI (100nM) and Multimeric-APRIL (100ng/ml - unless otherwise stated).

For gene expression experiments, cells were seeded at day 6 at 1 × 10^6^/ml in phenol red free media supplemented with 0.5% HIFBS, IL6 (10 ng/ml), IL21 (10 ng/ml) and GSI (100nM) for 16-20 hours. Multimeric-APRIL (100ng/ml) or SDF1 (1 ng/ml) was added and cells analyzed at indicated times.

For culture of ex vivo cells, following isolation cells were cultured in media containing IL6 (10ng/ml) and IL21 (10ng/ml) for 24 hours. Cells were then harvested and transferred into media containing IL6 (10ng/ml) and either Multimeric-APRIL (100ng/ml) and GSI (100nM); or IFNα (100u/ml) for 14 days. Half the media was replenished after 7 days.

### Flow cytometric analysis

Cells were analyzed using 4-to 6-color direct immunofluorescence staining on a Cytoflex LX or S (Beckman Coulter) flow cytometer. Antibodies used were: CD19 PE (LT19), CD138 APC (44F9) (Miltenyi); CD20 e450 (2H7) (eBioscience); CD27 FITC (M-T271), CD38 PECy7 (HB7), CD54 PE (HA58) (BD Biosciences). Controls were isotype-matched antibodies or FMOs. Dead cells were excluded by 7-AAD (BD Biosciences). Absolute cell counts were performed with CountBright beads (Invitrogen). Cell populations were gated on FSC and SSC profiles for viable cells determined independently in preliminary and parallel experiments. Analysis was performed with FlowJo version 10 (BD Biosciences) and Prism 8 (GraphPad). Statistical analysis performed was either Two-tailed paired T-test or RM one-way ANOVA, Tukey’s multiple comparisons test.

### RNA, cDNA and RT-PCR

RNA was extracted with TRIzol (Invitrogen), subjected to DNAseI treatment (DNAfree, Ambion) and reverse transcribed using Superscript II Reverse Transcriptase (Invitrogen). Taqman^®^ Assays for *FOS* (Hs00170630_m1), *FOSB* (Hs00171851_m1), *EGR1* (Hs00152928_m1) and *PPP6C* (Hs00254827_m1) were carried out according to manufacturer’s instructions and run on a Stratagene Mx3005p.

### Protein analysis

At the indicated time points, cells were lysed in Laemmli buffer. For cytoplasmic:nuclear protein samples, proteins were extracted using a cytoplasmic and nuclear extraction kit (Boster Bio) and protein concentration determined by BCA assay (Boster Bio). Samples were separated by SDS-PAGE and transferred to nitrocellulose. Proteins were detected by ECL (SuperSignal WestPico PLUS, Thermo Scienctific) and visualized on a ChemiDoc (BioRad) or film. Protein bands were quantitated using Image Lab 6.0.1 software (BioRad).

Antibodies used were p-AKT, AKT, p-ERK1/2, ERK1/2, p-FOXO1/3/4, FOXO1, FOXO3, p-JNK, JNK, p-p38, p38, MYC, RELA, p-SQSTM1 T269/S272, SQSTM1 (CST); TUBULIN (Merck); BLIMP-1 R23;(22) goat anti-mouse HRP, goat anti-rabbit HRP (Jackson ImmunoResearch).

### ELISpot

Influenza specific immunoglobulin was detected as previously described.(23) Inactivated influenza vaccine manufactured by Sanofi Pasteur MSD was used for coating plates.

For detection of human IgG secretion, a Human ELISpot^BASIC^ IgG kit (Mabtech) was used. The assay was performed as described in the manufacturer’s protocol, and 1000/2000 cells were added as indicated in the figure. Cells were incubated on plates for 16-20 hours in IMDM containing either standard amounts of IL6 and IL21 (control), or IL6 with either IFNα or APRIL/GSI.

### Gene expression data acquisition and analysis

Gene expression data sets were generated from differentiating PBs (day 7). At day 6, PBs from 4 healthy donors were seeded at 1 × 10^6^/ml in phenol red free IMDM supplemented with 0.5% HIFBS, IL6 (10 ng/ml), IL21 (10 ng/ml) and GSI (100nM) and incubated for 20 hours. Multimeric-APRIL (100ng/ml) was then added. A pre-treatment sample (0 minutes) and post treatment samples were removed at +30, +60, +120 and +360 minutes.

RNA was obtained using TRIzol (Invitrogen) and sequencing libraries generated with a TruSeq Stranded Total RNA Human/Mouse/Rat kit (Illumina). Libraries were sequenced on a NextSeq500 platform (Illumina), using 76-bp single-end sequencing, for details of fastq files initial quality assessment, trimming, alignment and annotation see supplemental methods. Transcript abundance was estimated using RSEM v1.3.0 and processed using DESeq2 to determine differential gene expression. Expression data sets are available with GEO accession GSE173644.

### Network analysis

For details of the Parsimonious Gene Correlation Network Analysis (PGCNA) approach see supplemental methods.(24) For the bulk RNA-seq network transcripts differentially expressed across the timeseries data (DESeq2 LRT FDR < 0.01) were merged per gene by taking the median value for transcript sets with a Pearson correlation ≥0.2 and the maximum value for those with a correlation <0.2 giving a 4,615 × 20 matrix. This was used for a PGCNA2 analysis (-n 1000, -b 100) giving a network with 16 modules. The median expression per timepoint was visualized as Z-scores mapped onto the network. The top 25 genes per module by network strength were used to generate Module Expression Values by summing their Z-scores (normalized across cells) per cell and visualized as a hierarchically clustered heatmap.

### scRNA-seq networks

The Croote et al peripheral blood single cell data was downloaded as counts per gene (Table SI of manuscript), consisting of 973 cells.(25) Cells with biased expression of genes related to growth-factor-signaling/IEG likely to reflect a handling artefact were identified and filtered (see Supplemental Methods). Data for the 655 remaining cells we re-filtered to retain consistently expressed genes (count ≥ 5 in ≥ 10% of cells) yielding 3,436 genes. PGCNA was carried out generating a network with 22 modules. The genes per module were summed and the transposed matrix used to generate a PGCNA network of the cells.

### Network availability

Interactive networks and meta-data are available at https://mcare.link/STC-APRIL.

### Gene signature data and enrichment analysis

A set of 40,686 signatures was generated by merging gene-ontology and gene-signatures as previously described (see Supplemental Methods).(24) Enrichment of gene lists for signatures was assessed using a hypergeometric test, in which the draw is the gene list genes, the successes are the signature genes, and the population is the genes present on the platform.

### Heatmap visualizations

The gene expression data and GSE results were both visualized using the Broad GENE-E package (https://software.broadinstitute.org/GENE-E/). For visualization of expression data, the Module Expression Values were visualized on a global scale. For GSE the signatures were filtered (FDR <0.1 and ≥ 5 and ≤ 1500 genes for the signature sets, selecting the top 15 most significant signatures per module) and the enrichment/depletion z-scores visualized. In both cases the data was hierarchically clustered (Pearson correlations and average linkage).

### Ethical approval

Approval for this study was provided by UK National Research Ethics Service via the Leeds East Research Ethics Committee, approval reference: 07/Q1206/47.

## Results

### APRIL supports in vitro PC survival which is enhanced by γ-secretase inhibition

*We* have previously defined conditions which allowed the generation of long-lived PCs in vitro both using stromal support and independent of stroma using type-1 IFN or TGFB mediated survival signals.(5, 6) These conditions allowed PC survival in the absence of defined TNFSF signaling, and in the absence of detectable NFκB mediated transcriptional response as assessed by gene signature analysis.(3, 5) This was notable since TNFSF members and in particular APRIL are considered to act at least in part through the provision of an NFκB pathway signal,(7) and APRIL can support PC survival in vitro.(26, 27) We therefore aimed to analyse a model system in which an NFκB signal was delivered as the PC completed differentiation.

We first used the ability of APRIL to support PC survival as a functional indicator of effective signaling. Initially performing a dose response, we observed that PC survival could be effectively supported by APRIL in multimeric form but required significant quantities (Figure 1A). Under these conditions the phenotype of the differentiated cells was consistent with an early PC state showing strong CD38 expression, partial upregulation of CD138 and loss of CD20 (Figure 1B). Recently it was observed that BCMA, the primary surface receptor for APRIL, expressed on ASCs was subject to active proteolytic shedding and that this was dependent on γ-secretase activity.(10) We therefore tested whether this effect was observed in ASCs in the model system. Indeed, γ-secretase inhibition substantially augmented the expression of cell surface BCMA during in vitro differentiation. A 6-fold enhancement of BCMA expression was observed following γ-secretase inhibition (Figure 1C), and this increase was largely maintained in the presence of APRIL stimulation (Figure 1D).

**Figure 1.**
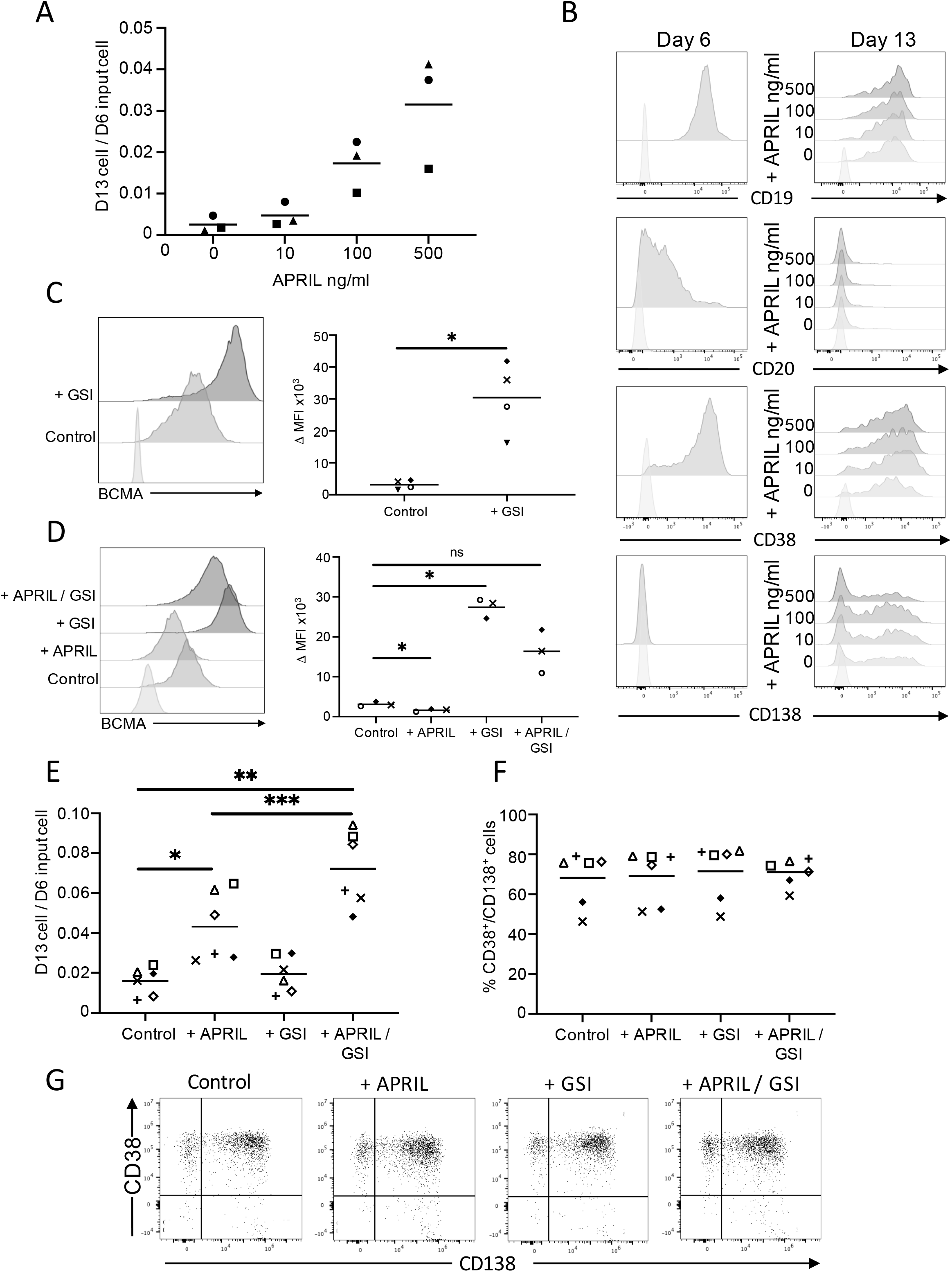
APRIL and γ-secretase inhibition support in vitro PC differentiation and survival. **(A)** APRIL dose response showing recovered cell number at day 13 of in vitro culture, y-axis - fraction of cells recovered at day 13 (D13) per input cell at day 6 (D6), x-axis - concentration of multimeric APRIL (ng/ml) from day 6 (all conditions included a standard dose of IL6 and IL21), data are shown for 3 donors (symbols). **(B)** Representative flow cytometry plots for selected antigens. Left panel - phenotype at day 6, before the addition of APRIL. Right panel - phenotype at day 13 following culture in IL6, IL21 and APRIL. Results are shown for each APRIL concentration equivalent to part (A) and antigens highlighted above each panel. **(C)** Impact of γ-secretase inhibition (GSI) on BCMA expression. Left panel - representative flow cytometry data for surface BCMA expression_at day 7 following treatment with indicated conditions. Right panel −ΔMFI (×10^3^) for BCMA expression against isotype control. Data are shown-for 4 donors (paired t-test: * p<0.05). **(D)** Impact of APRIL treatment on BCMA expression in the presence or absence of GSI. Left panel − representative flow cytometry data for surface BCMA expression_at day 13 following treatment with indicated conditions. Right panel −ΔMFI (×10^3^) for BCMA expression against isotype control. Data shown are for 3 donors (RM one-way ANOVA test: * p<0.05). **(E)** Cell number recovered after APRIL stimulation from day 6 to day 13 of in vitro culture under conditions indicated (x-axis), y-axis - fraction of cells recovered at day 13 (D13) per input cell at day 6 (D6) (RM one-way ANOVA test: * p<0.05, ** p<0.01, *** p<0.001). **(F)** Percentage of CD38^+^/CD138^+^ cells observed at day 13 in samples cultured as in (E). **(G)** Representative scatter plots of CD38 vs CD138 expression of a single donor at day 13, with culture conditions indicated above each panel. Data shown in E & F are representative of 6 donors. In parts C-E, control conditions are media plus standard dose of IL6 and IL21. Individual donors are indicated by unique symbols which are consistent across all figures.

The enhancement of surface BCMA expression following γ-secretase inhibition translated into a significant increase in the impact of APRIL on in vitro PC survival (Figure 1E). This increase in viability was associated with a generally similar phenotype of cell populations (Figure 1F and G). Thus, the APRIL mediated survival benefit for PC populations in vitro can be enhanced by inhibition of BCMA shedding in a fashion consistent with the model proposed by Laurent et al.(10) and providing further evidence that surface shedding is an intrinsic feature limiting BCMA signals at the PB to PC transition.

### APRIL support for ex vivo PB survival is enhanced by γ-secretase inhibition

Since the combination of APRIL and γ-secretase inhibition provided an effective condition for in vitro derived PB/PC transition we next sought to determine whether this would also provide support for ex vivo PBs. We therefore isolated PBs from 5 donors following seasonal influenza vaccination at day-7 of the vaccine response and transferred these cells into survival conditions with either IFNα or APRIL and γ-secretase inhibition. Two weeks later we assessed the phenotype, number and secretory function of the PC population (Figure 2). For all donors tested, the number of viable cells at two weeks was significantly greater in the presence of APRIL and γ-secretase inhibition than in the presence of IFNα (Figure 2A). Phenotypically the conditions were similar in generating CD19^lo^ CD27^hi^ CD38^hi^ and CD138^hi^ PCs (Figure 2B). However, consistently in the presence of APRIL and γ-secretase inhibition the expression level of CD19 and CD27 was higher and the CD38 expression lower (Figure 2C). Functionally the cell populations generated were indistinguishable at the level of per cell secretion of IgG as assessed by ELIspot and included influenza vaccine specific ASCs (Figure 2D and E and Supplemental Figure 1 B and C). We conclude therefore that IFNα and APRIL, which provide distinct signals, can each promote survival and maturation of ex vivo PBs sustaining a similar population of PCs in terms of antibody secretion and phenotype. The subtle but reproducible differences in surface phenotype observed for in vitro differentiated PCs under distinct niche conditions supports the contention that intracellular signals activated by the survival niche in which a PB matures impact on the functional state of the resulting PC.

**Figure 2.**
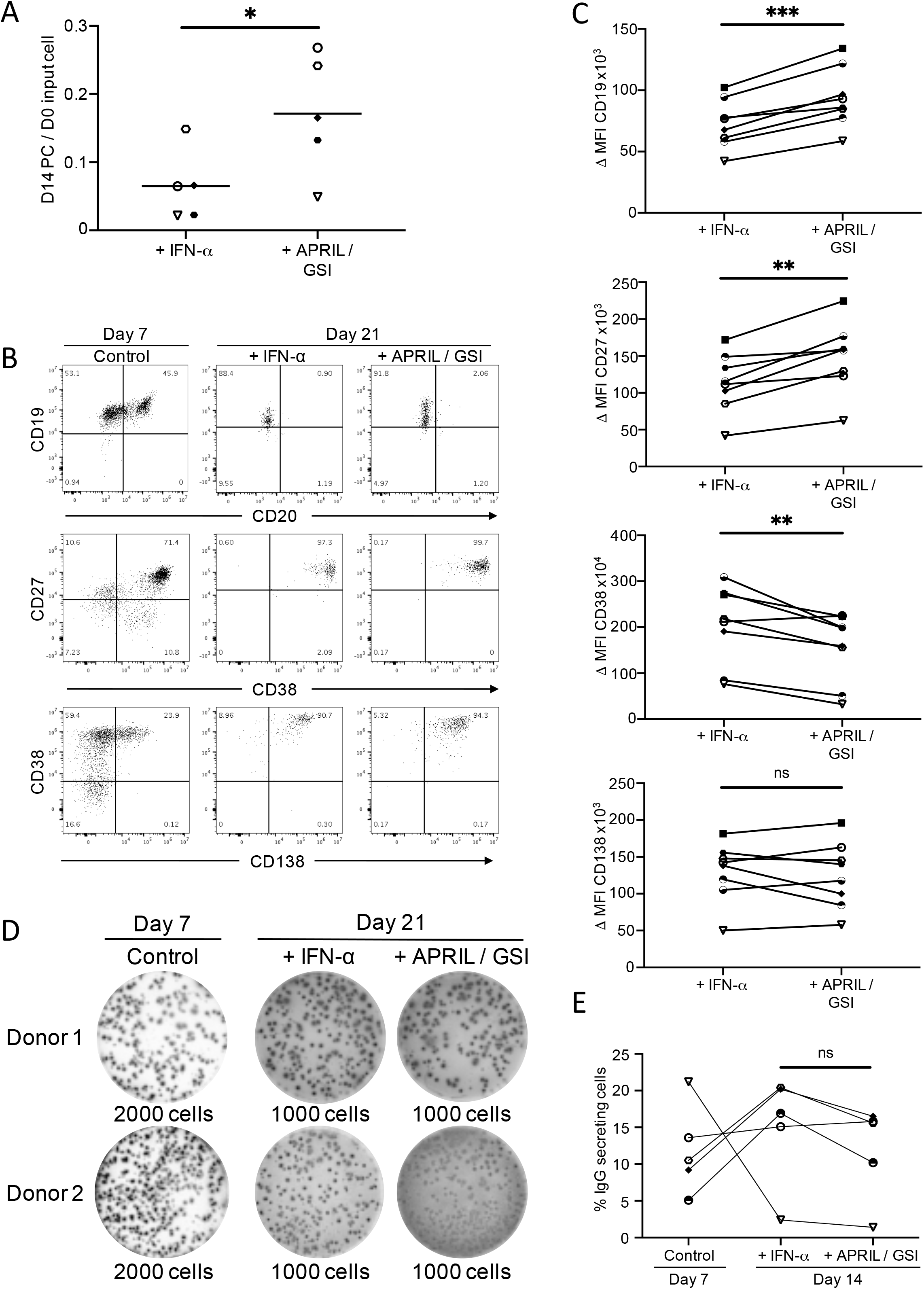
APRIL supports ex vivo PB maturation. **(A)** Comparison of cell recovery after 14 days of in vitro culture for ex vivo BCMA+ PBs isolated at day 7 after influenza vaccination cultured in IL6 and either IFNα or APRIL/GSI conditions (x-axis), y-axis – PCs at day 14 (D14) per input cell at day 0 (D0) (paired t-test * p<0.05). Data shown are for 5 donors (symbols). **(B)** Representative phenotypes of cells isolated ex vivo at day 7 after influenza vaccination left panels, or after 14 days of in vitro culture (equivalent to day 21 post vaccination) in IL6 with IFNα (middle) or APRIL/GSI (right) panels. Scatter plots from top to bottom show CD19/CD20, CD27/CD38 and CD38/CD138 as indicated. **(C)** Differential expression for antigens assessed in (B), shown in order CD19, CD27, CD38 and CD138 from top to bottom as ΔMFI (×10^3^ or ×10^4^ as indicated) against isotype control (y-axis) for IFNα (left) or APRIL/GSI (right) (x-axis). Each donor is identified with a unique symbol (paired t-test: ns - not significant, ** p<0.01, *** p<0.001). Data shown are for 8 donors. **(D and E)** Equivalent immunoglobulin secretion is supported by either IFNα or APRIL/GSI conditions. Representative ELISpot results for two independent donors from ex vivo isolated cells at day 7 post influenza vaccination (left panels) or after 14 days of in vitro culture with IL6 and either IFNα (middle) or APRIL/GSI (right) panels, equivalent to day 21 post influenza vaccination (D) and quantitation (E) shown for 5 donors.

### APRIL drives sustained activation of MAP kinase pathways

Having established conditions under which APRIL supported survival of both in vitro generated and ex vivo derived PBs and allowed maturation of these populations to the PC state, we were in a position to evaluate the downstream signaling pathways regulated during this response. We focused on the initial transition when the PB encounters the APRIL signal. We have previously shown that another PC niche signal, SDF1, drives potent activation of ERK MAP-kinase within 5 minutes of activation.(6) We therefore initially evaluated whether this was a response shared with APRIL. In contrast to SDF1 stimulation, APRIL induced a more delayed activation of ERK peaking at 30 minutes (Figure 3A). APRIL also activated p38 and JNK, which was sustained for 120 minutes following APRIL stimulation (Figure 3B and C). We tested whether the differences in upstream pathway activation might translate into differences in immediate early gene (IEG) regulation and indeed APRIL showed both a more modest amplitude for *FOS, FOSB* and *EGR1* induction, as well as a more delayed kinetics of peak response for *FOS* and *EGR1* compared to SDF1 (Fig. 3E). Thus, APRIL drives activation of MAP-kinase pathway in PBs and leads to induction of IEGs, with different kinetics from that observed with SDF1.

**Figure 3.**
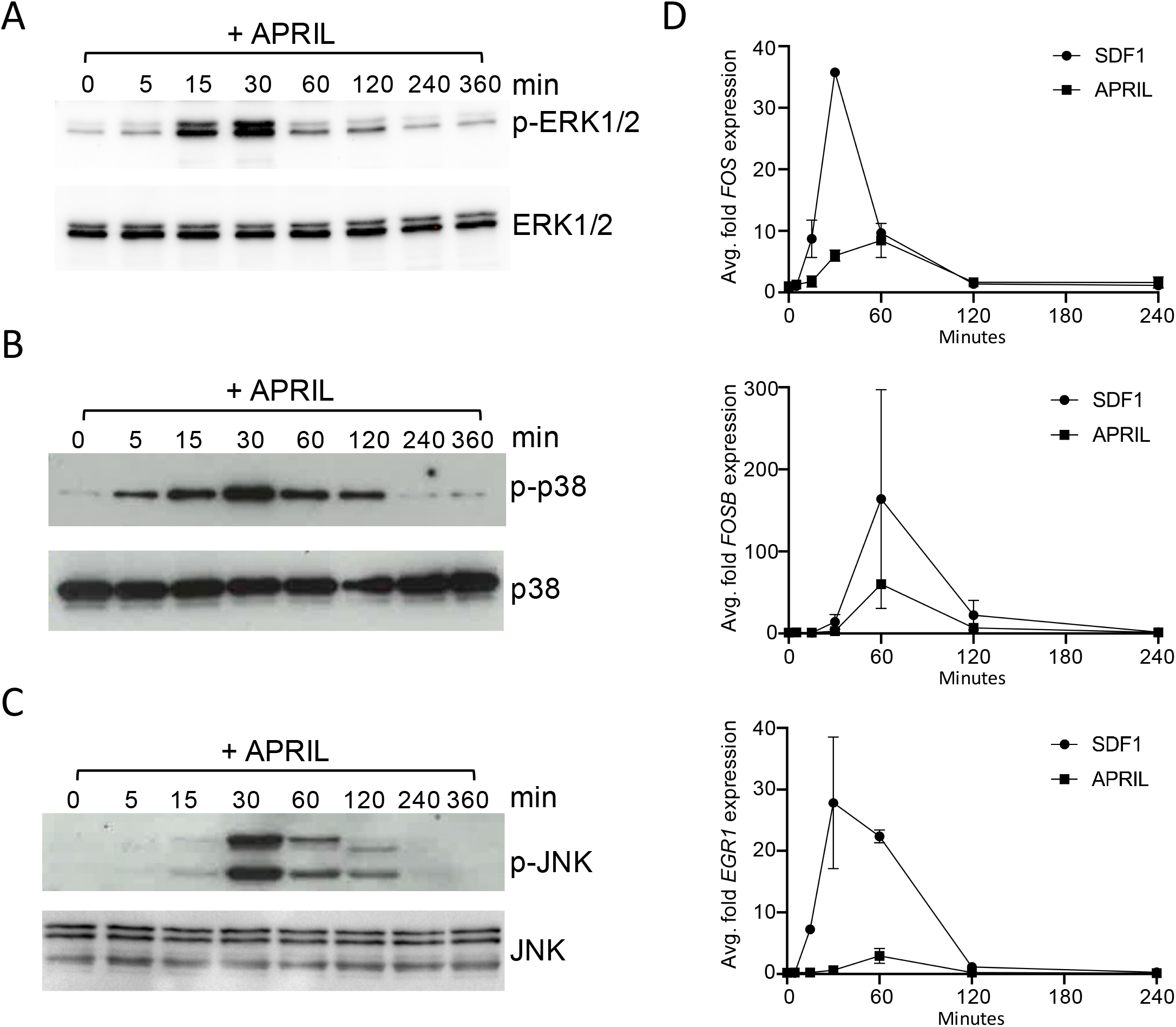
MAPkinase pathway activation and immediate early gene regulation in response to APRIL. **(A)** Time course of ERK1/2 phosphorylation induced after stimulation of day 7 PBs with APRIL. Upper panel show detection of phospho-ERK1/2 (p-ERK) after stimulation with APRIL (left lanes) for the indicated time points of t= 0 (unstimulated), 5, 15, 30, 60, 120, 240 and 360 minutes. Total ERK1/2 loading control is shown below. **(B)** Time course of p38 phosphorylation induced by APRIL. Upper panel show detection of phospho-p38 (p-p38) after stimulation with APRIL for the time course as in (A). Total p38 loading control is shown below. **(C)** Time course of JNK phosphorylation induced by APRIL. Upper panel show detection of phospho-JNK1/2 (p-JNK) after stimulation with APRIL for the time course as in (A). Total JNK loading control is shown below. Western blots shown in (A-C) are representative of 6 donors. **(D)** Relative kinetics of immediate early gene induction following stimulation of day 7 PBs with SDF1 (filled circles) or APRIL (filled squares). Expression of *FOS, FOSB* and *EGR1* is shown in order from top to bottom as average fold expression relative to t=0 normalized against housekeeping control and detected by qRT-PCR from RNA isolated at t= 0 (unstimulated), 5, 15, 30, 60, 120, 240 and 360 minutes after stimulation. Data are shown as average and standard deviation of n=2 (SDF1) and n=4 (APRIL) donors.

### APRIL activates classical NFκB responses and induces AKT phosphorylation and FOXO1 nuclear exclusion

The NFκB pathway is considered to be of central importance for activation and survival signaling downstream of TNF receptor super family (TNFRSF) members and can itself drive expression of IEGs. In cell lines APRIL has been principally linked to activation of the classical NFκB pathway.(18) Consistent with this we observed rapid induction of IκBα phosphorylation and subsequent sustained loss of IκBα protein following APRIL stimulation of PBs (Figure 4A). This phosphorylation and loss of IκBα also correlated with nuclear translocation of RELA (Figure 4B). While CD40L stimulation also induced IκBα phosphorylation and subsequent sustained loss of IκBα protein in PBs (Figure 4A), unlike APRIL, CD40L addition at the PB stage contributed to subsequent expansion of cells with retained B-cell features, rather than promoting the emergence of a phenotypic PC population (Supplemental Figure 2 A and B). Thus, these TNFSF/TNFRSF pairs have quite distinct impacts on the fate of differentiating ASC populations when encountered at the PB stage with APRIL promoting both NFκB activation and supporting PC differentiation.

**Figure 4.**
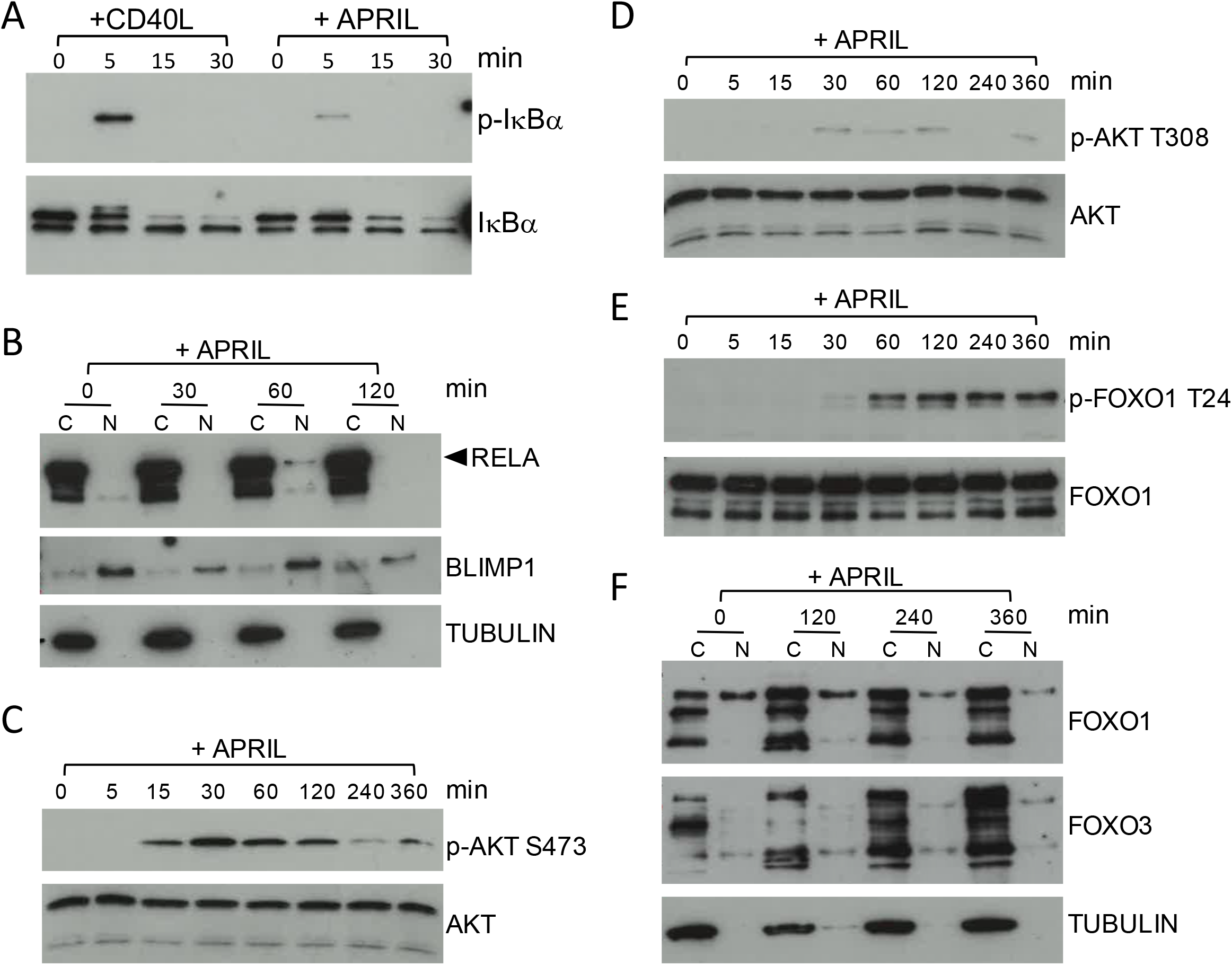
Activation of NFκB/RELA and AKT/FOXO1 pathway by APRIL stimulation. **(A)** Time course of IκBα phosphorylation after simulation of day 7 PBs with APRIL compared to CD40L. Upper panel show detection of phospho-IκBα (p-IκBα) after stimulation with CD40L (left lanes) or APRIL (right lanes) for the indicated time points of t=0 (unstimulated) 5, 15 and 30 minutes. Total IκBα is shown below. Stimulation leads to loss of IκBα at later time points. Data representative of 4 donors. **(B)** Nuclear localisation of RELA following APRIL stimulation. PBs at day 7 were unstimulated (t=0) or stimulated with APRIL 30, 60 and 120 minutes and cytoplasmic and nuclear fractions were separated as indicated above the lanes (C and N). Samples were immunoblotted for RELA (arrow, upper panel), BLIMP1 (nuclear enriched, middle panel) and TUBULIN (cytoplasmic fraction, lower panel). Data representative of 4 donors. **(C, D** and **E)** Activation of AKT/FOXO1 pathway after APRIL stimulation. Day-7 PB were unstimulated (t=0) or stimulated with APRIL for 5, 15, 30, 60, 120, 240 and 360 minutes. Samples were probed for **(C)** AKT phospho-serine 473 (p-AKT S473) and total AKT; and **(D)** AKT phospho-threonine 308 (p-AKT T308) and total AKT; and **(E)** FOXO1 phospho-threonine 24 (p-FOXO1 T24) and total FOXO1 as indicated. Data representative of 4 donors (C & D), 5 donors (E). **(F)** Cytoplasmic and nuclear extracts (C and N above the lanes) from PBs at day 7 were unstimulated (t=0) or treated with APRIL for 120, 240 or 360 minutes were blotted for FOXO1 (upper panel), FOXO3 (middle panel) and TUBULIN (lower panel). Data representative of 4 donors.

PI3kinase pathway activation leading to AKT phosphorylation and regulation of FOXO family members is a critical element of survival and activation signaling during earlier stages of B-cell differentiation. Recently it has been proposed that stromal mediated survival signals may contribute specifically to PC survival through activation of the AKT-FOXO pathway, while APRIL acts more selectively via NFκB. We therefore examined whether APRIL signaling could induce AKT activation.(27) Indeed, induced phosphorylation of AKT at both S473 and T308 was observed in response to APRIL (Figure 4C and D). This showed a prolonged kinetics and was maintained to 120 minutes after stimulation. Although S473 phosphorylation was more intense than that of T308, full activation of the pathway was supported by induced phosphorylation of FOXO1/3 with a kinetics consistent with the pattern of AKT activation (Figure 4E). Both FOXO1 and FOXO3 were expressed in PBs, but FOXO3 was restricted to the cytoplasmic fraction prior to stimulation. In contrast FOXO1 was present in both cytoplasmic and nuclear fractions and showed evidence of nuclear exclusion at later time points after APRIL treatment, consistent with an active signaling in response (Figure 4F). The kinetics of this response paralleled that of other acute signaling pathways, and in the context of the model tested was independent of stromal contact. Therefore, APRIL signals can suffice to activate the AKT-FOXO pathway and drive FOXO1 nuclear exclusion at the PB to PC transition.

### APRIL signals propagate to a robust gene expression response

We next sought to evaluate the overall impact of APRIL on gene expression in differentiating PBs. We applied a combination of gene expression time course and parsimonious gene correlation network analysis (PGCNA),(28) evaluating samples with RNAseq at 30, 60, 120 and 360 minutes after stimulation with APRIL. We analyzed the data for differentially expressed genes across the time series using a likelihood ratio test (FDR <0.01; Supplemental Table 1 available online). The resulting 4,615 genes were used to generate a gene correlation network which resolved into 16 modules (Figure 5A, Supplemental Tables 2 and 3 available online; https://mcare.link/STC-APRIL). We analysed the biology associated with these gene expression modules using gene signature and ontology enrichment analysis (Supplemental Figure 3 and online resources). This demonstrated a distinct segregation of enriched biology across the early response gene modules (M2 and M3) and secondary response gene modules (M6, M9 and M13). For each module a suitable summary term was derived from the observed enriched ontologies and signatures.

**Figure 5.**
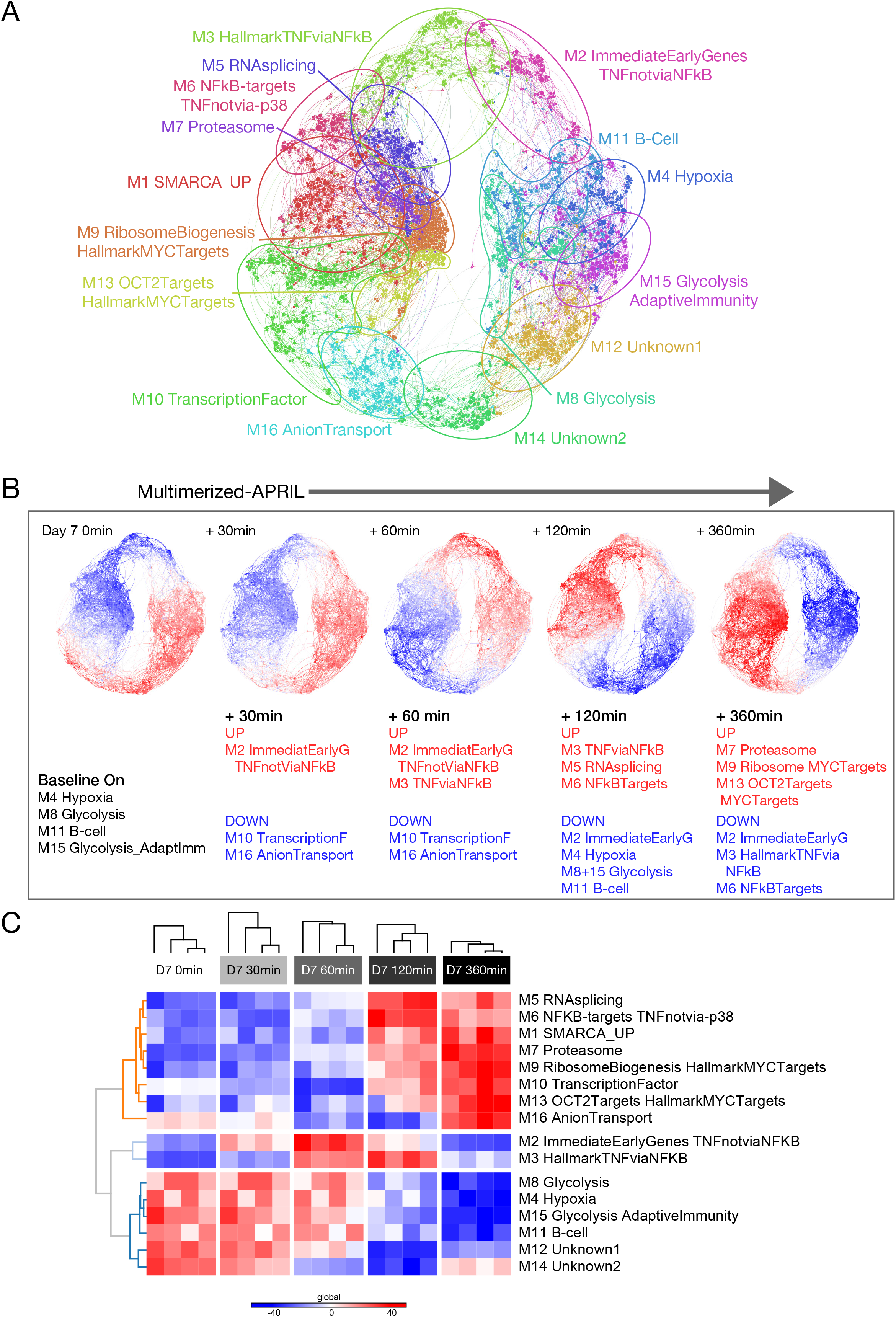
The transcriptional response of PBs to APRIL stimulation. **(A)** PGCNA network representation of the modular pattern of gene expression induced following APRIL stimulation of day 7 PBs over a 360 minutes time course. Network modules M1 to M16 are colour coded and are designated with a summary term derived from gene ontology and signature separation between network modules. For interactive version go to https://mcare.link/STC-APRIL. Visualization of top gene ontology and signature enrichments between network modules are provided in Supplemental Figure 3 and lists of module genes and ontology enrichments in Supplemental Tables 2 and 3 available online. **(B)** Overlay of gene expression z-scores for all genes in the network shown in blue (low) to red (high) Z-score color scale across the time course indicated by the arrow above the panel from left to right. Left panel unstimulated t=0 followed by expression patterns at t= 30, 60, 120 and 240 minutes from left to right as indicated in the figure. Beneath each colour coded network select upregulated and down regulated modules at each time point are indicated using module number and summary term (red-upregulated module expression, blue-downregulated module expression). **(C)** Summary representation of patterns of expression across all network modules as a heat map shown as module expression values derived from the top 25 genes of each module as Z-scores with a colour scale (blue (−50 low) red (+50 high)). Samples and modules are separated by hierarchical clustering. Module numbers and indicative module terms are shown on the right.

To assess the kinetics of gene expression change, the relative gene expression was overlaid on the network and assessed as a Module Expression Value (MEV) heatmap (Figure 5B and C and online resources). This revealed a distinct wave of gene expression propagating around the network. The initial activation was observed in module M2, enriched for IEGs, and genes linked to the gene signature of “TNF response not mediated by NFκB”. This is consistent with an initial wave of gene expression downstream of MAP kinase pathway activation, which was followed at 60 minutes by the induction of modules enriched for genes linked to NFκB signaling and the response to TNF (M3). This module (M3) was sustained to 120 minutes at which time it was joined by a wider diversity of NFκB target genes (M6) and a module of genes enriched for factors involved in RNA splicing. Finally, by 360 minutes as the initial signaling modules (M2, M3 and M6) waned in expression the secondary response of gene expression was enriched for modules related to MYC (M9 and M13) and OCT2 targets (M13) along with the ribosome (M9) and proteasome (M7). Thus, the response to APRIL follows a classical pattern involving a dominant initial impact on IEGs followed by NFκB response modules and ultimately leads to the expression of functional gene modules that indicate a contribution for MYC and OCT2 in gene regulation. Indeed, while PBs express modest amounts of MYC relative to activated B-cells, MYC protein remained detectable across the time course and showed modest induction following APRIL stimulation by 60 to 120 minutes (Supplemental Figure 2C-E).

### Expression of MCL1 correlates principally with ASC state rather than APRIL response

A mechanism that may couple APRIL/BCMA signaling to PC survival is the suppression of apoptosis through regulation of MCL1.(15) While we found modest evidence of upregulation for *MCL1* and *BCL2* following APRIL stimulation of PBs, the response for *MCL1* in particular was subtle and occurred in the context of significant *MCL1* expression at all stages of the time course. This suggested that expression of *MCL1* was a feature of the PB state largely independent of APRIL stimulation. To assess whether this was also a feature of primary human ex vivo ASCs at the PB stage we turned to an external data source, taking advantage of single cell gene expression data which has been generated from peripheral blood B-cells and PBs.(25) To analyse these data in an analogous fashion to our gene expression time course, we modified the PGCNA approach and applied this to the single cell expression data. The resulting network resolved into 22 modules (Figure 6A, Supplemental Tables 4 and 5 available online) separating into modules representative of the B-cell state (sc_M3) (Supplemental Figure 4 and online resources), and primary features of the PC state (sc_M10). Other modules separated expression features related to B-cell activation and PC differentiation (sc_M1), exosomes and adhesion (sc_M2), ribosomes/translation (sc_M5), proliferation/MYC-target genes (sc_M7) and mitochondrial genes (sc_M22). The resolved modules of gene expression were differentially expressed between individual cells and allowed separation of clusters of resting or activated B-cells from PBs (Figure 6B).

**Figure 6.**
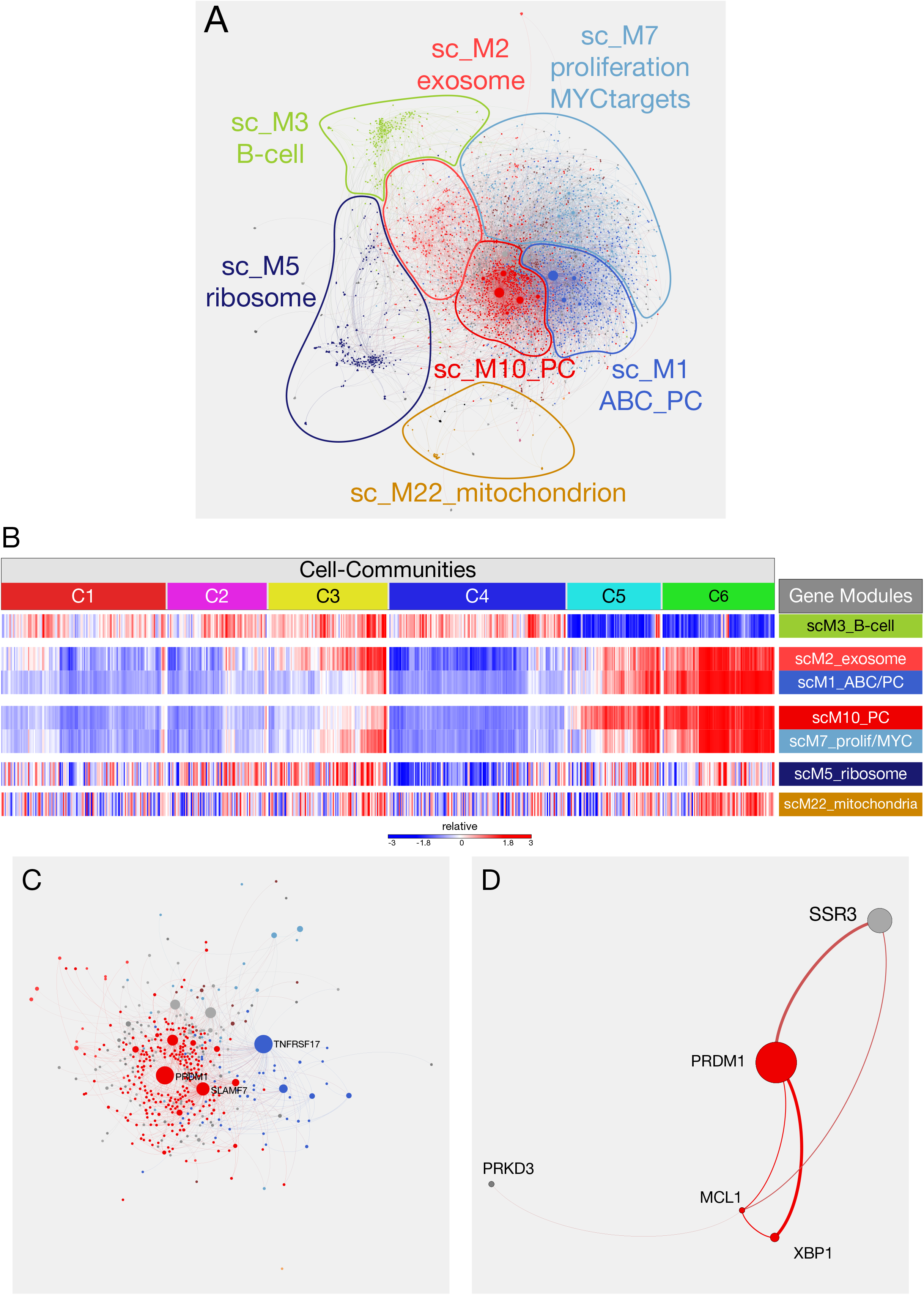
Network visualisation of single cell gene expression in human peripheral blood PBs/PCs. **(A)** Single cell gene expression data of human peripheral blood B-cells and PB/PCs were imported from Croote et al. (25) and analyzed with PGCNA to generate a network visualization of gene expression in single cell gene expression comparable to analysis to bulk expression data. Genes are identified by nodes in the network and retained direct correlations by edges. Node sizes are determined by betweenness-centraliy. Network modules are color-coded and location within the network identified with matching outlines and indicative module names. For interactive version go to https://mcare.link/STC-APRIL. Visualization of top gene ontology and signature enrichments between network modules are provided in Supplemental Figure 4 and lists of module genes and ontology enrichments in Supplemental Tables 4 and 5 available online. **(B)** Clustering of cells with adaptation of PGCNA. Clusters of cells (cell-communities) were derived by carrying out a parsimonious correlation analysis on the transposed matrix used in part (A) giving 6 (C1-C6) communities. Cells within each community were subsequently hierarchically clustered based on summed gene module expression values (Pearson correlations and average linkage). Summed expression values for each cell for the modules identified in (A) and mentioned in the text are shown as a heat map with indicative gene module names identified on the right. Cellcommunities C1-C6 are identified above. **(C)** *PRDM1* as the hub gene of PC expression module sc_M10. The panel highlights the neighbours of *PRDM1* in the PGCNA network. Genes represented by nodes in the network are colour coded according to module membership as in (A), node size is determined by betweenness-centrality. Gene names are shown for connected hub genes (SLAMF7 (sc_M10 red) and TNRFSF17 (sc_M1_ABC/PC royal blue)). (D) The nearest neighbours in the network for *MCL1*, which is not a hub gene, but includes *PRDM1* and *XBP1* amongst its four neighbours.

*PRDM1*, encoding BLIMP1 a master regulator of PC differentiation, was identified as the hub gene of module sc_M10 encompassing features of the PC state. PGCNA is based on initial radical edge reduction for all correlated genes (n=3 connections per gene retained), hub nodes emerge by virtue of being amongst the most correlated partners of many other genes. As a hub gene, *PRDM1* was highly interconnected. Its immediate neighbours included other core genes of the PC state, such as *IRF4, XBP1* and *SLAMF7(CD319)* (Figure 6C). *MCL1* was also an immediate neighbour of *PRDM1. MCL1* itself was not a hub gene in the network but its immediate most correlated gene neighbours included both *PRDM1* and *XBP1* (Figure 6D). Thus, the expression of *MCL1* correlates with the key transcriptional regulators of the PB/PC state in ex vivo peripheral blood PBs. We conclude therefore that expression of *MCL1* in PBs before APRIL exposure is consistent with the physiological state of peripheral blood PBs prior to entry into survival niche conditions.

### The APRIL response includes regulation of cell adhesion and metabolism genes

A dominant feature of the APRIL response lies in the regulation of the NFκB pathway, and amongst the most significantly induced genes were several related to adhesion and metabolism including *ICAM1, NINJ1, NAMPT* and *SQSTM1*. In fact, these genes also belong to the Hallmark signature of TNFA signaling via NFκB and are thus consistent with a canonical response to TNFSF/TNFRSF interaction. *ICAM1* and *NINJ1* are both involved in the process of cell adhesion, and ICAM1 interactions via stromal fibroblasts have been recently highlighted as synergising with APRIL signals to support PC survival via PI3K signaling and FOXO phosphorylation.(27) While APRIL signaling was independently able to sustain activation of this pathway in our model, regulation of ICAM1 would nonetheless provide the potential for sustained signaling. We therefore examined ICAM1 expression following APRIL stimulation by flow cytometry, consistent with the gene expression data ICAM1 expression was significantly upregulated on the surface of PCs within 24h of APRIL stimulation (Figure 7A and B).

**Figure 7.**
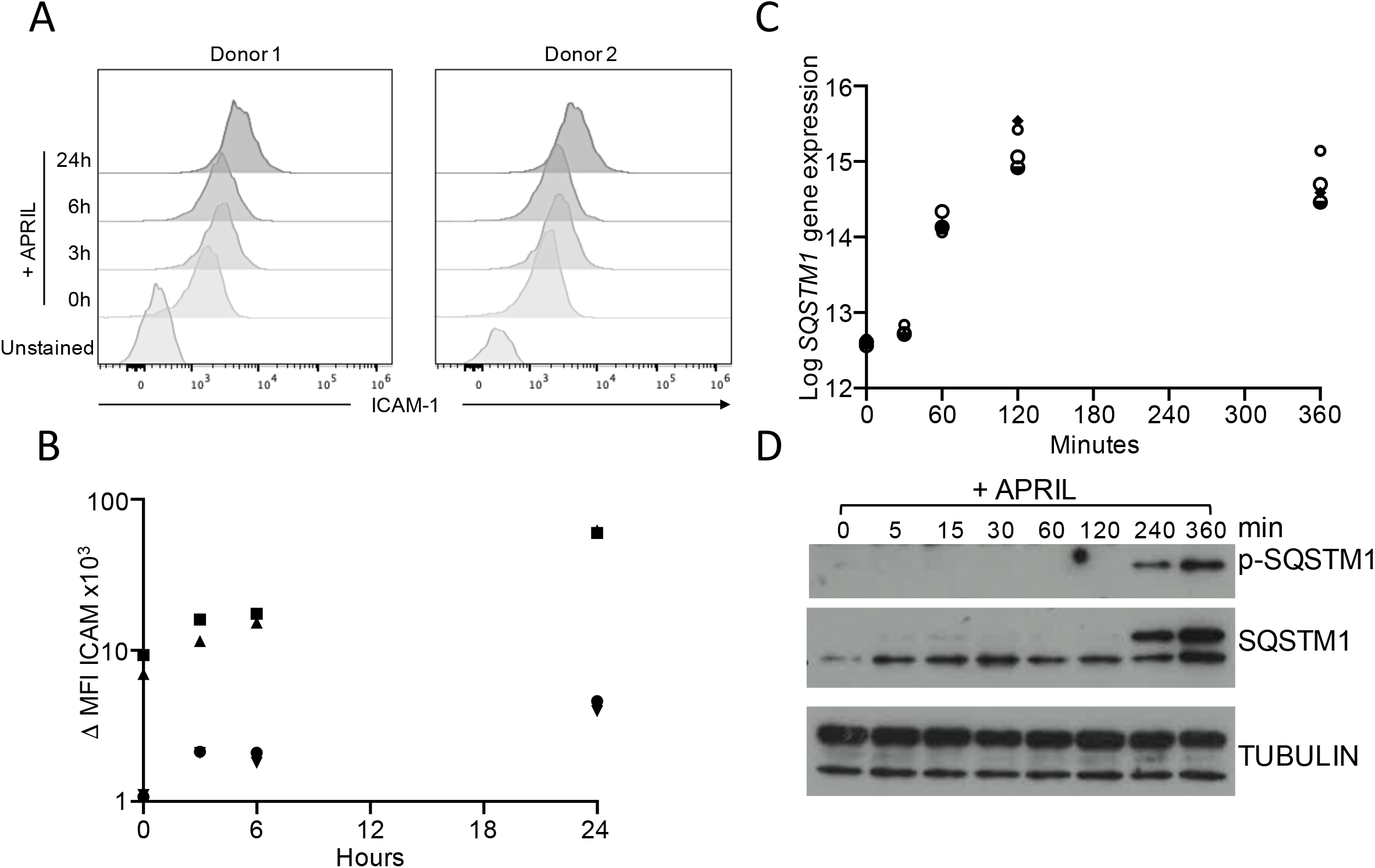
ICAM1 and SQSTM1 expression are induced in response to APRIL in PBs. **(A)** Expression of ICAM1 was detected by flow cytometry in day 7 PBs after APRIL stimulation. Upper panels show histograms of ICAM1 expression (x-axis) against unstained controls for 2 representative donors with lanes from bottom of each panel in the order unstained control, t = 0 and at 3h, 6h and 24h after APRIL stimulation. **(B)** Shows ΔMFI (×10^3^) of ICAM1 expression against unstained control in 4 donors identified with symbols. **(C)** Expression of *SQSTM1* induced following APRIL stimulation with log expression values derived from RNA-seq expression profiling as in Figure 5, with individual donors shown with symbols. **(D)** Shows a representative Western blot of n=4 for SQSTM1 expression after APRIL stimulation, unstimulated (t=0) or stimulated with APRIL for 5, 15, 30, 60, 120, 240 and 360 minutes shown above the lanes. Upper panel: phospho-SQSTM1; middle panel: total SQSTM1; lower panel: TUBULIN loading control.

Metabolic regulation and autophagy are important factors in PC longevity, and *SQSTM1* expression is both a feature of the PB/PC state and induced following APRIL exposure. SQSTM1 is a multifunctional protein with roles in selective autophagy, metabolic regulation via mTORC1 and regulation of the NFκB pathway. We found that SQSTM1 was both robustly induced in expression and was phosphorylated in differentiating PBs, following APRIL stimulation. Phosphorylation of SQSTM1 was observed with antibody specific to Thr269/Serine272 a target site for p38,(29) which suggests that SQSTM1 may provide a point of integration for APRIL induced signals in PBs (Figure 7C and D). Thus, the control of genes linked to the canonical TNFSF/TNRSF response pathway along with the downstream regulation of modules related to MYC and OCT2 point to a diverse impact of APRIL on PB/PC biology which together provide several pathways through which survival may be sustained.

## Discussion

The generation and maintenance of long-lived PCs from transitional PB populations is dependent in vivo on the migration of a PB to micro-anatomical niches that provide suitable signals for survival.(4) It is postulated that once localized to such a niche the PC must retain the capacity to reside in and respond to the niche signals provided, or be displaced by newly generated ASCs. Such competition potentially limits PC longevity, and both extrinsic niche factors and intrinsic features of the differentiating ASC may contribute to determine competitive fitness.(1) However understanding how niche signals impact on the maturation of PCs and the signaling pathways employed has been limited by intrinsic features of PC biology including limited cell number and inaccessible anatomical location in particular in the context of human PCs.

Amongst the niche signals that have been defined as important for in vivo long lived PC generation in vivo are signals delivered by the TNFSF member APRIL.(30) Here we have used an in vitro model system to address the signaling and gene regulatory response in human PBs induced by encounter with APRIL. Our data demonstrate that the acute signaling responses are diverse, including activation of canonical NFκB, p38 and JNK MAPkinase as well as AKT. This pattern shows considerable overlap to signaling responses induced by BAFF in B-cells and recapitulates the pattern of signaling demonstrated for BCMA activation in the context of forced over-expression in cell line models.(7, 18) This confirms that a broad diversity of signaling is induced by this factor in differentiating primary human ASCs. APRIL also activates the ERK MAPkinase pathway but with relatively delayed kinetics when compared to the response to the chemokine SDF1 (CXCL12).(6) Such differences in upstream signalling also propagate into distinct kinetics of IEG response, and downstream transcriptional responses. In a reductionist model of niche homing the response to the chemoattractant SDF1 would precede responses to signals linked to niche residence such as APRIL. Niche size can be understood in terms of signaling microdomains with diffusion and consumption combining to limit the range over which particular niche signals may act.(31) Combining temporal and spatial sequences of niche signals with differential downstream pathway activation provides the potential to encode complex gene regulatory responses which may in turn contribute to differences in functional specialization and fitness of individual PCs.

The precise pathways responsible for signal propagation at the plasma membrane following exposure to APRIL remain to be determined. We confirmed the observation of Laurent et al. that γ-secretase inhibition maximises surface BCMA expression.(10) This increase in BCMA expression translated into enhanced APRIL induced survival responses. In conjunction with the existing literature it is therefore most likely that the dominant receptor for the APRIL response in this model is indeed BCMA. While TACI expression is not substantially impacted by γ-secretase treatment it is expressed at low levels in PBs, and thus a contribution to signal propagation cannot be excluded (data not shown). Future studies will be needed to address the membrane proximal signaling events that lead to the activation of the diverse downstream pathways. Regulation of AKT phosphorylation and by inference activation of PI3K is of particular interest. We note for example that APRIL signaling preferentially maintains CD19 expression amongst the differentiating PCs, and that CD19 has been identified as a hub for PI3K activation in the B-lineage.(32)

Our analysis shows that the APRIL induced signaling pathway activation propagates into successive waves of gene regulation. These follow a classical pattern of immediate early, delayed early and secondary response gene regulation.(33) The modules of genes that correspond to immediate and delayed early responses show a high degree of overlap to patterns of gene regulation identified in the TNFA signaling hallmark gene sets (Broad GSEA).(34) The most immediate responses are the control of co-ordinated modules of genes shared with the TNF response that have been attributed to NFκB independent signaling, and most likely relate to MAPkinase pathway induction. These are closely followed by typical NFκB response modules which include many of the negative feedback regulators typically associated with canonical NFκB pathway activation and including miR genes such as *MIR3142HG*, which is the host gene for miR-146a, a negative regulator of NFκB pathway activation.(35) An essential role for miR-155 in sustaining murine PB proliferation and class-switched antibody production has been recently reported,(36) and the induction of *MIR155HG* as human PBs mature in response to APRIL suggests that an important role for this miRNA continues into the quiescent PC state.

As the gene regulatory response to APRIL propagates, the initial immediate early and NFκB target genes are repressed, consistent with efficient negative feedback regulation of the response. At the same time the propagation of the signal into secondary response genes focuses in particular on ribosome, MYC and OCT2 related gene signatures. The secondary response modules induced by APRIL do include selected elements related to direct control of cell cycle for example *CCND2* and *CDK4*, but enrichment of cell cycle related signatures in large part reflects coordinated induction of multiple proteasome subunit genes. This argues that while APRIL signals do impact on cell cycle related gene expression, the secondary response modules are principally related to ribosome function, RNA processing and biogenesis and hence to cell growth rather than cell proliferation per se.

In mice *MCL1* has been shown to be essential for PC survival, and regulated independently of BLIMP1 during PC differentiation.(15) In human PBs, whether in vitro generated or in vivo derived, we find that *MCL1* mRNA expression is closely correlated with core features of the PC state. In murine PCs expression of MCL1 is reduced in bone marrow but not splenic PC after genetic deletion of *BCMA*.(15) In keeping with this, we find that APRIL enhances the expression of both *BCL2* and *MCL1* in human PBs. However, as *MCL1* is already abundantly expressed its further induction is only modest over the time course tested. The in vitro data are consistent with the pattern of *MCL1* expression in single cell transcriptomic data of peripheral blood B-cells and PBs in which *MCL1* correlates with other core features of the PC state. As our analyses is focused on the earliest response of PBs to APRIL the data do not exclude a greater role in maintaining expression of MCL1 as long-lived PCs are subsequently established. However, it is likely that additional pathways contribute to the survival advantage conferred by APRIL during the initial PB/PC window. Amongst the most substantially induced genes over the time course are transcription factors such as *ATF3*, adhesion molecule *ICAM1*, and metabolism and autophagy related genes such as *NAMPT* and *SQSTM1*.

In a recent study it has been argued that long-lived (memory) PC survival depends on two key pathways, activation of NFκB via APRIL/BCMA and activation of the PI3K/AKT/FOXO pathway in response to integrin binding to ICAM1/VCAM1 on stromal cells.(27) Human PCs both in vitro generated and ex vivo derived have the potential for survival without contact dependent help from stromal cells, although such stromal cells may provide additional support.(5) Here we find that APRIL itself can induce phosphorylation of AKT consistent with PI3K pathway activation independent of a stromal cell contact. Additionally, amongst the most significantly induced genes following APRIL signaling in human PBs is *ICAM1* and corresponds with enhanced surface ICAM1 expression upon differentiation. This provides the potential means to support homotypic adhesion. Cohesive masses of PCs are a characteristic feature in inflammation and in PC neoplasia, for example in plasmacytomas,(37) arguing that homotypic adhesion and signaling can act as contributors to PC survival in vivo. Induction of ICAM1 by APRIL or downstream of related NFκB pathway activation provides one means through which such signaling may substitute for a stromal contact dependent pathway. However, while this may provide one mechanism the diverse signaling and gene regulator response induced by APRIL argue that multiple pathways are likely in sum to contribute to the prosurvival signal delivered by this niche factor at the PB to PC transition.

## Acknowledgements

This work was supported by Cancer Research UK program grant (C7845/A17723 and C7845/A29212), and Cancer Research accelerator award (C355/A26819).

We thank Ulf Klein for critical review of the manuscript.

---

**Supplemental Figure 1 (accompanies Figure 2). APRIL supports ex vivo PB maturation. (A)** Representative phenotypes of cells isolated ex vivo at day 7 after influenza vaccination left panels, or after 14 days of in vitro culture (equivalent to day 21 post vaccination) with IL6 and either IFNα (middle) or APRIL/GSI (right) panels. Scatter plots from top to bottom show CD19/CD20, CD27/CD38 and CD38/CD138 as indicated for 4 individual donors. **(B and C)** Representative ELISpots for influenza specific ASCs shown for cells isolated at day 7 post vaccine response, or after 14 days of in vitro culture with IL6 and either IFNα (middle) or APRIL/GSI (right) panels, equivalent to day 21 post influenza vaccination (B) and quantitation (C). Numbers of cell seeded per well are shown below. Cells were incubated on plates for 16-20 hours in IMDM containing either standard amounts of IL6 and IL21 (Control, D7) or IL6 with either IFNα or APRIL/GSI (D21).

**Supplemental Figure 2 (accompanies Figure 4 and 5). Comparison of CD40 and APRIL from day 6 of culture. (A)** Representative phenotypes for cells from two donors at day 13 of culture with additional of APRIL/GSI (left panels) or soluble CD40L/GSI (right panels) along with supportive cytokines IL6 and IL21. Shown are scatter plots for expression of CD38 (y-axis) against CD138 (x-axis). **(B)** Upper panel recovered cell number at day 13 for PBs cultured from day 6 with APRIL/GSI (left) or soluble CD40L/GSI (right) (x-axis), y-axis displays day 13 (D13) cells as fraction of day 6 (D6) input. Lower panel percentage of CD38^+^/CD138^+^ cells at day 13 as percentage of viable cells (y-axis) for PBs cultured from day 6 with APRIL/GSI (left) or soluble CD40L/GSI (right) (x-axis) along with supportive cytokines IL6 and IL21. Four individual donors identified by symbols. **(C)** Expression of *MYC* mRNA induced following APRIL stimulation with log expression values derived from RNA-seq expression profiling as in Figure 5, with individual donors shown with symbols at the indicated time points in min (x-axis). **(D)** Representative Western blot of n=4 for MYC expression in day-7 PBs after APRIL stimulation, unstimulated (t=0) or stimulated with APRIL for 5, 15, 30, 60, 120, 240 and 360 minutes shown above the lanes. Upper panel: MYC; lower panel: TUBULIN loading control. **(E)** MYC protein expression quantified against TUBULIN loading control across an APRIL response time course as shown in (D) for 4 individual donors identified with unique symbols. Expression is normalized to 100% for all samples based on expression for each donor at t=0.

**Supplemental Figure 3 (accompanies Figure 5). Gene ontology and signature enrichments for gene modules of the PB response to APRIL.** Heatmap of gene ontology and signature term enrichments linked to the PGCNA modules of the time course network analysis for APRIL response (signatures were pre-filtered to p-value <0.001 and ≥ 5 and ≤ 1000 genes; selecting the top 15 most enriched signatures per module). For full signature enrichment lists, please see Supplemental Table 3. Modules are shown along the x-axis, and signature terms along the y-axis. Signature terms and modules are hierarchically clustered to illustrate relationships. Enrichment (red) and depletion (blue) of signatures are shown on colour scale of z-score.

**Supplemental Figure 4 (accompanies Figure 6). Gene ontology and signature enrichments for gene modules of the single ASC/B-cell expression network derived from Croote et al. data set.** Heatmap of gene ontology and signature term enrichments linked to the PGCNA modules derived from single cell gene expression in Figure 6(A) (signatures were pre-filtered to p-value <0.001 and ≥ 5 and ≤ 1000 genes; selecting the top 15 most enriched signatures per module). For full signature enrichment lists, please see Supplemental Table 5. Modules are shown along the x-axis, and signature terms along the y-axis. Signature terms and modules are hierarchically clustered to illustrate relationships. Enrichment (red) and depletion (blue) of signatures are shown on colour scale of z-score.

