## Supplementary figures and images for "APRIL drives a co-ordinated but diverse response as a foundation for plasma cell longevity"

### Supplemental Figure 1

#### Slide 1
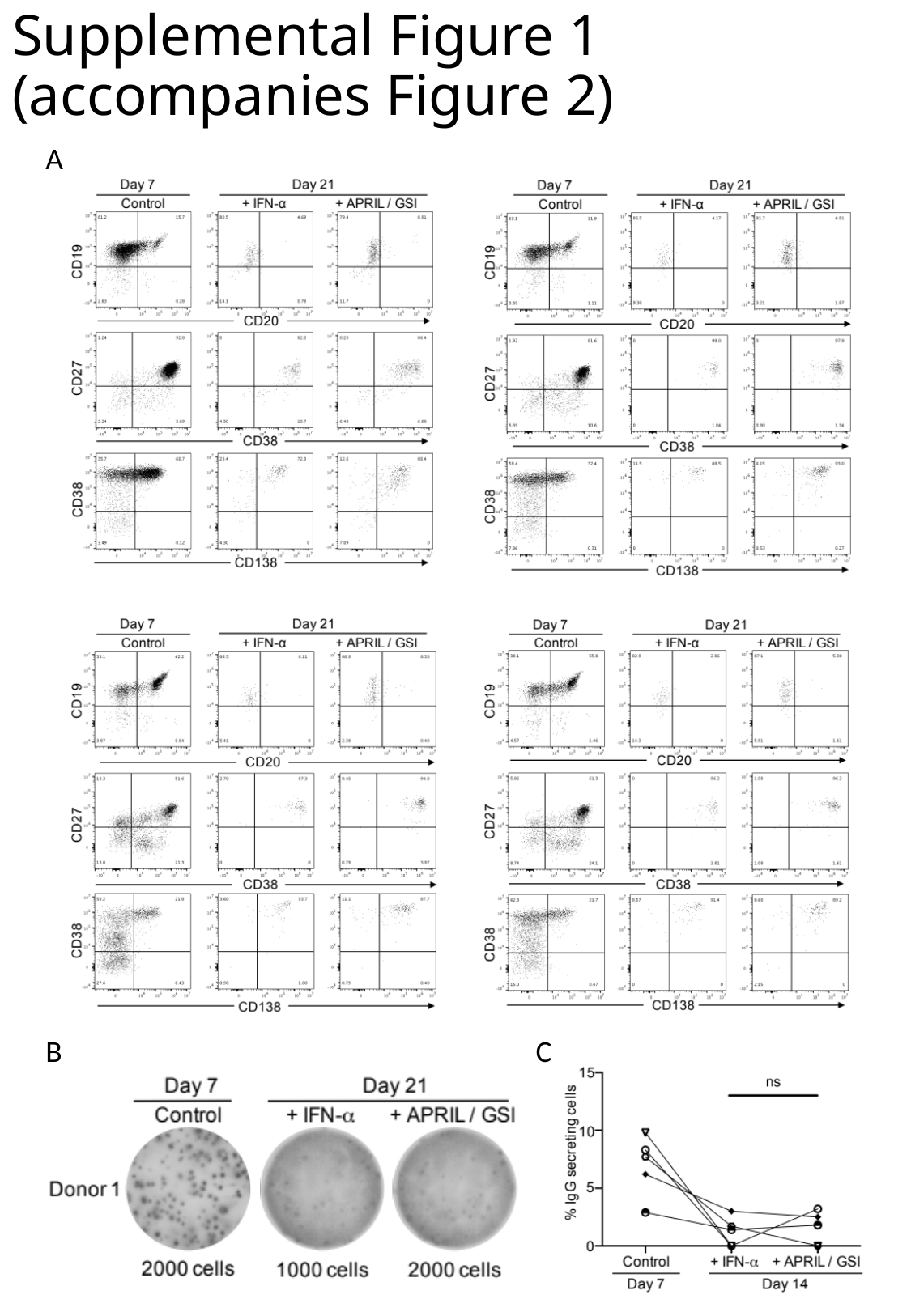

##### Supplemental Figure 1 (accompanies Figure 2)
A
B
C

### Supplemental Figure 2

#### Slide 1
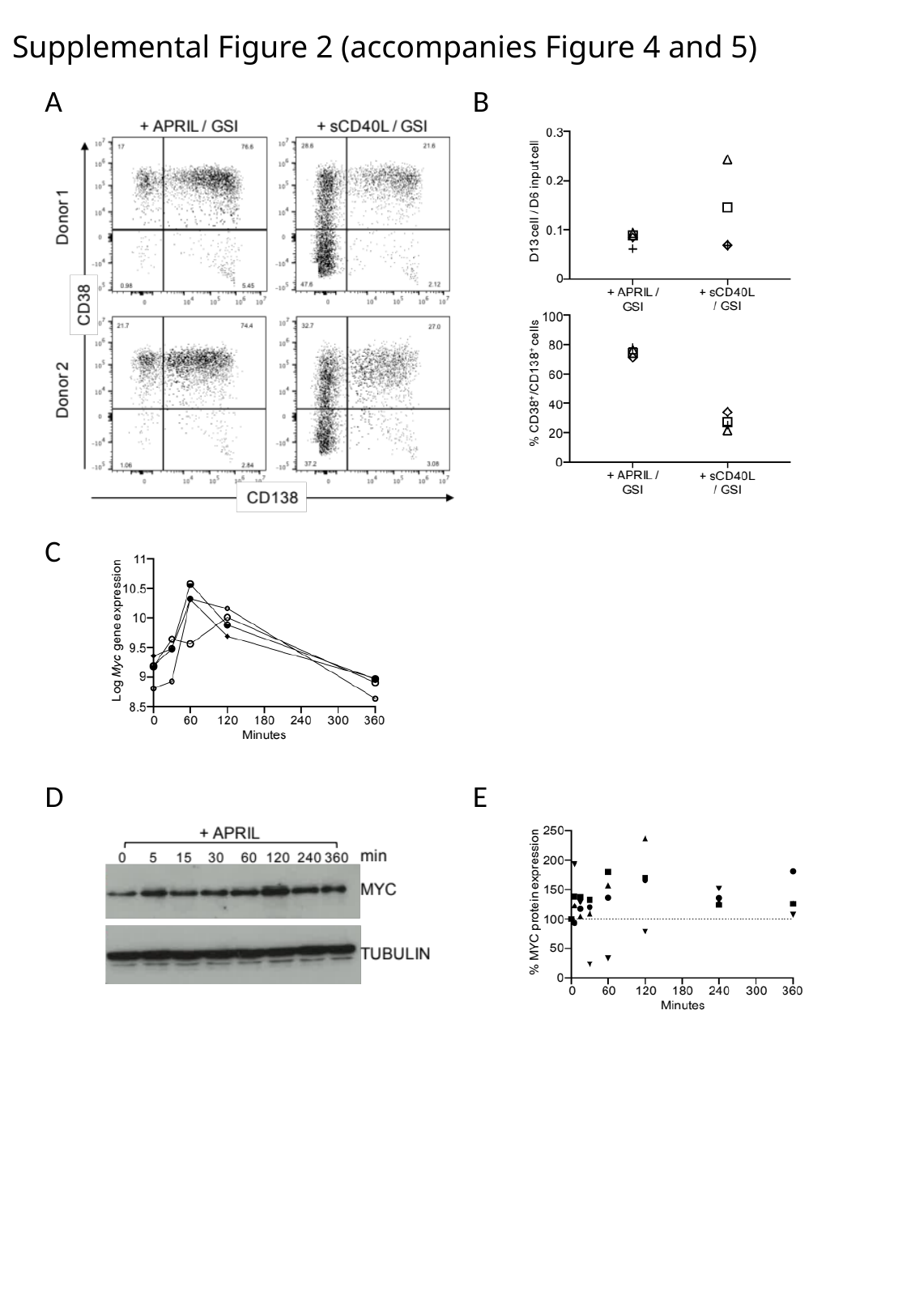

##### Supplemental Figure 2 (accompanies Figure 4 and 5)
A
B
C
D
E

### Supplemental Figure 3

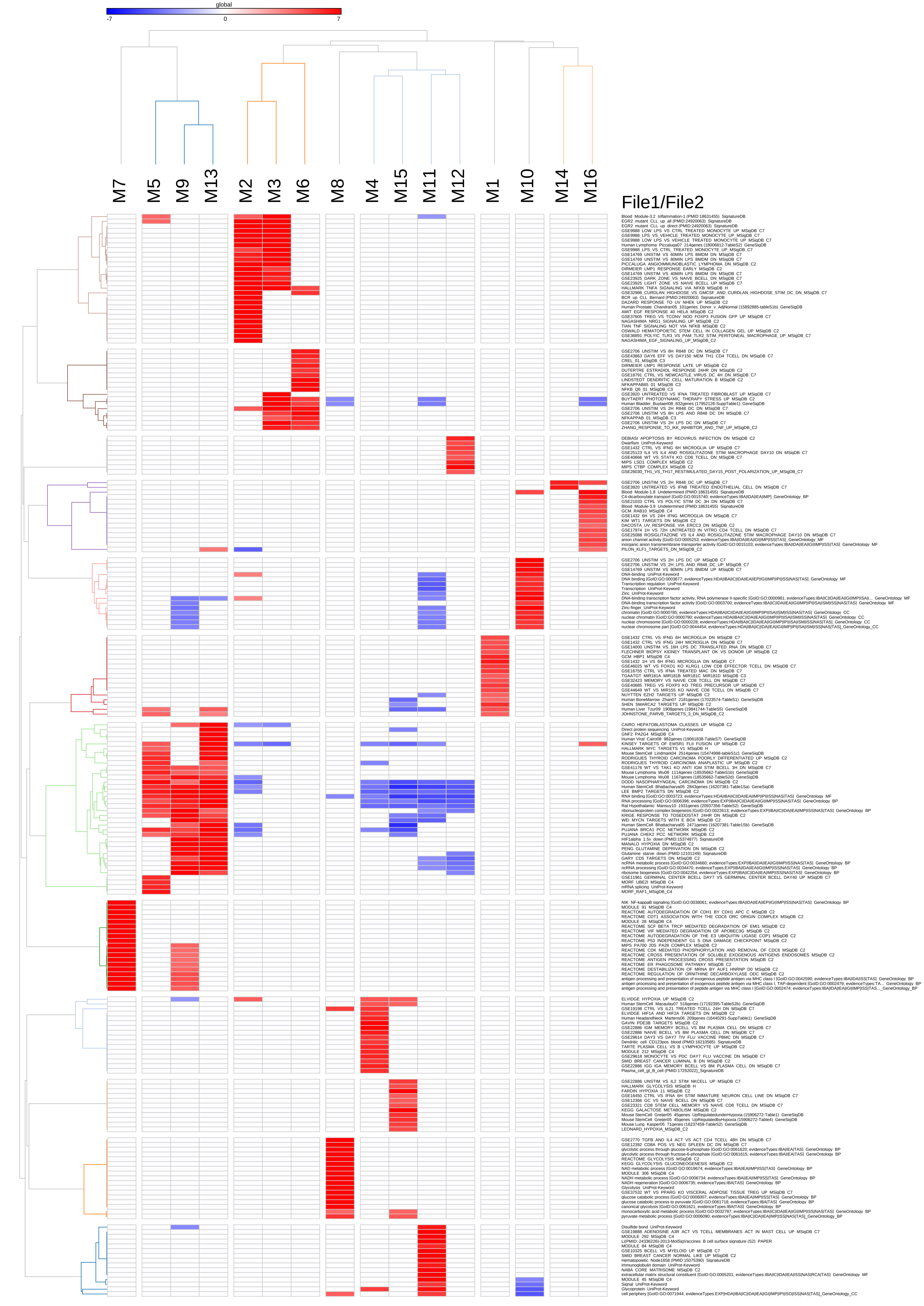

### Supplemental Figure 4

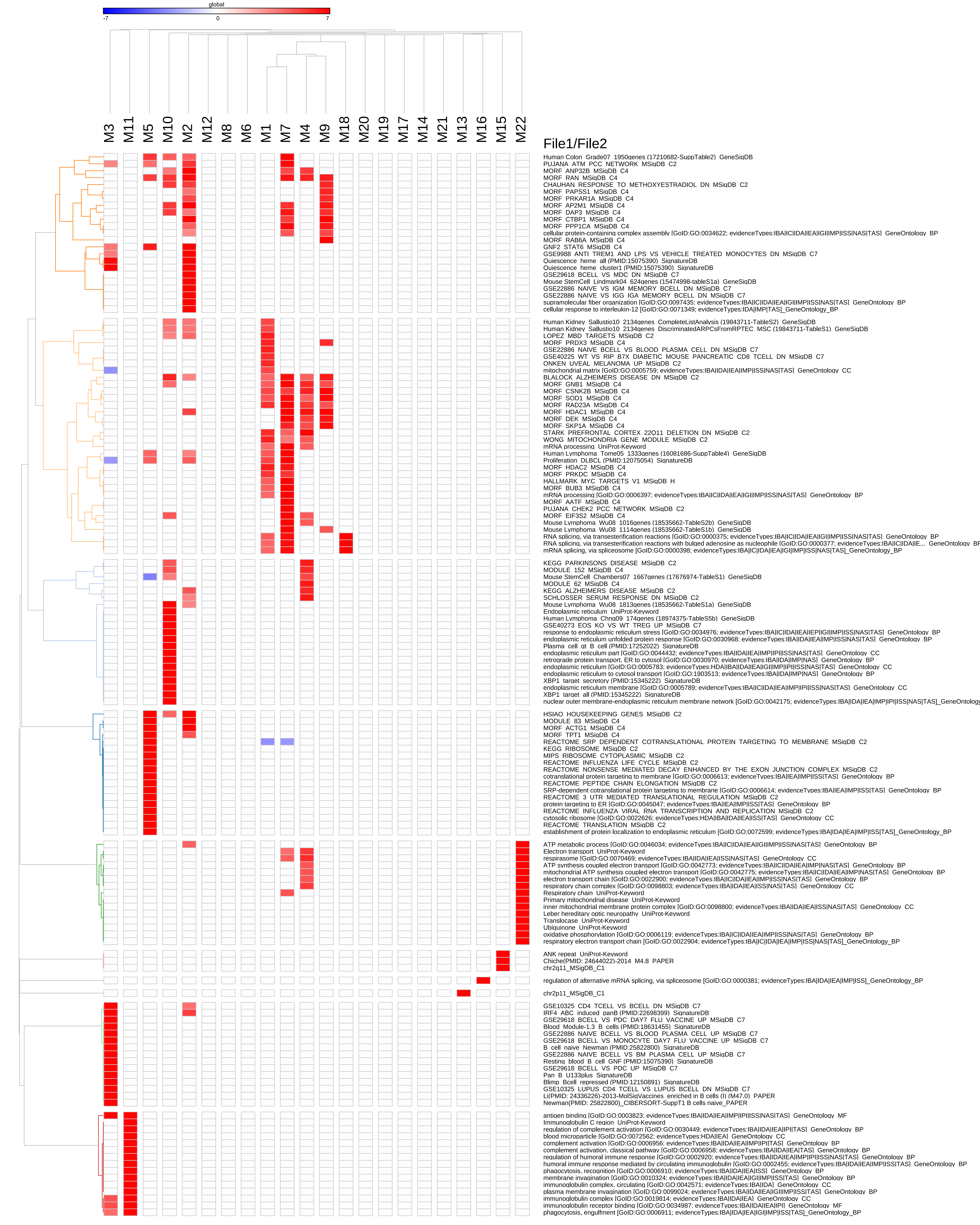
