## Supplemental Methods for "APRIL drives a co-ordinated but diverse response as a foundation for plasma cell longevity"

### RNA-seq analysis

#### Expression data sets

Gene expression data was generated from differentiating PBs (day 7) from 4 healthy donors maintained in low serum media with IL6/IL21 and then post treatment with APRIL at + 30 , 60, 120 and 360 minutes.

#### Analysis

RNA was obtained using TRIzol (Invtirogen) and sequencing libraries generated with a TruSeq Stranded Total RNA Human/Mouse/Rat kit (Illumina). Libraries were sequenced on a NextSeq500 platform (Illumina), using 76-bp single-end sequencing. The fastq files were assessed for initial quality using FastQC v0.11.8 (https://www.bioinformatics.babraham.ac.uk/projects/fastqc/), trimmed for adapter sequences using TrimGalore v0.6.0 (https://www.bioinformatics.babraham.ac.uk/projects/trim_galore/) and aligned to GRCh38.p12/hg38 (Gencode release 28) using STAR aligner (v2.6.0c) using twopassMode[1]. Transcripts were re-annotated using the MyGene.info (http://mygene.info) API using all available references and any ambiguous mappings manually assigned. Transcript abundance was estimated using RSEM v1.3.0 and imported into R v3.5.1 using txImport v1.10.1 and then processed using DESeq2 v1.22.2[2–5]. Using DESeq2 differential gene expression was determined (using LRT) quality visualised using MA plots and shrinkage of log fold estimated using the apeglm method (Supplemental Table 1).[6]

### PGCNA networks

#### Background

For details and validation of the Parsimonious Gene Correlation Network Analysis (PGCNA) approach see our other work.[7] Here a brief description of the method will be given. After informative genes are selected they are used to calculate Spearman's rank correlations for all gene pairs. For each gene (row) in the correlation matrix only the 3 most correlated edges per gene are retained. The resulting matrix $M$, with entries written as $M = (m_{ij})$ is made symmetrical by setting $m_{ij} = m_{ji}$ for all indices $i$ and $j$ so that $M = \mathrm{M}^{\mathrm{T}}$ (its transpose). The resultant correlation matrix is clustered using a community detection algorithm (Leidenalg v0.7.0) and the best (judged by modularity score) used for downstream analysis. For the networks generated here we carried out 1000 clusterings and analysed the top 100 for biological enrichment against a signature database (see Gene signature data) by generating Scaled cluster enrichment scores (see original PGCNA methods for details).[7,8]

#### Bulk RNA-seq network

The transcripts differentially expressed across the timeseries data (DESeq2 LRT FDR < 0.01) were merged per gene by taking the median value for transcript sets with a Pearson correlation ≥0.2 and the maximum value for those with a correlation <0.2 giving a 4,615 x 20 matrix. This was used for a PGCNA2 analysis (-n 1000, -b 100) giving a network with 16 modules (Supplemental Table 2, Figure 5A). The median expression per timepoint was visualised as Z-scores mapped onto the network (Figure 5B). For each gene in the network a strength (edge-weight x degree) was calculated and used to select the top 25 genes per module. These were then converted to Module Expression Values by summing their Z-scores (normalised across cells) per cell and visualised as a hierarchically clustered heatmap (Figure 5C).

#### scRNA-seq networks

The Croote *et al* peripheral blood single cell data was downloaded as counts per gene (Table S1 of manuscript), consisting of 973 cells.[9] The data was filtered to only retain the highly expressed genes (count ≥ 5 in ≥ 10% of cells) yielding 4,349 genes. This matrix was used for PGCNA analysis (using Pearson correlation; -n 1000, -b 100), generating a network with 19 modules. The genes per module were summed for each cell and visualised as a heatmap, revealing a subset of cells with almost sole expression of M10 (data not shown). M10 is enriched for growth-factor-signalling/IEG, which we believe in this context represents a dissociation/handling artefact. Thus, we removed these samples (n=318) and for the remaining 655 cells we re-filtered the total set of genes to retain the highly expressed genes (count ≥ 5 in ≥ 10% of cells) yielding 3,436 genes. Another PGCNA analysis was carried out generating a network with 22 modules (Figure 6A). The same matrix was then transposed and used to generate a PGCNA network (using Pearson correlation; -n 1000, -b 100) of the cell-communities. In this case the best clustering of the data was selected based on the Leidenalg modularity score. The genes per module (from part A) were summed and then hierarchically clustered (Pearson correlations and average linkage) within each cell-community group discovered in the PGCNA cell network (Figure 6B).

### Data visualisation

#### Network visualisation

The optimal PGCNA networks were converted to a list of edges and nodes and uploaded into the Gephi package (version 0.9.2).[10] Degree and Betweenness Centrality were calculated, and the latter used to adjust node sizes. The network layout was generated using the ForceAtlas2 approach, and interactive HTML5 web visualizations exported using the sigma.js library (https://github.com/oxfordinternetinstitute/gephi-plugins/tree/sigmaexporter-plugin). The interactive visualisations can be found at https://mcare.link/STC-APRIL.

#### Heatmaps

The gene expression data and GSE results were both visualised using the Broad GENE-E package (https://software.broadinstitute.org/GENE-E/). For visualisation of expression data, the Module Expression Values (see Bulk RNA-seq network) were visualised on a global scale. For GSE the signatures were filtered (FDR <0.1 and ≥ 5 and ≤ 1500 genes for the signature sets: selecting the top 15 most significant signatures per module) and the enrichment/depletion z-scores visualised. In both cases the data was hierarchically clustered (Pearson correlations and average linkage).

### Statistical analyses

#### Gene signature data

A dataset of 17,904 gene signatures was created by merging signatures downloaded from http://lymphochip.nih.gov/signaturedb/ (SignatureDB), http://www.broadinstitute.org/gsea/msigdb/index.jsp MSigDB V6.2 (MSigDB C1–C7 and H; excluding C5. With MIPS signatures from version 3.1 and PID signatures from version 4 added back), http://compbio.dfci.harvard.edu/genesigdb/ Gene Signature Database V4 (GeneSigDB), UniProt keywords (parsed XML from http://www.uniprot.org/downloads), and fifteen papers.[11–28] A gene ontology gene set was created using an in-house python script. This parses a gene association file (http://geneontology.org/page/download-go-annotations) to link genes with ontology terms and then uses the ontology structure (.obo file; http://purl.obolibrary.org/obo/go.obo) to propagate these terms up to the root. The resultant gene set contained 22,782 terms. The gene-ontology and gene-signatures sets were merged to give a final signature set of 40,686 terms.

#### Enrichment analysis

The gene signature enrichment (GSE) was assessed using a hypergeometric test, in which the draw is the gene list genes, the successes are the signature genes, and the population is the genes present on the platform. The resultant p-values are then adjusted for multiple testing using Benjamini and Hochberg correction. For the PGCNA networks GSE analyses the genes per module were compared against the 40,686-signature database (background: all the genes in that network).

### Data processing

All analyses were undertaken on ARC3, part of the High Performance Computing facilities at the University of Leeds, UK.

### Data and software availability

Interactive networks and all meta-data are available at https://mcare.link/STC-APRIL. PGCNA python scripts are available at https://github.com/medmaca/PGCNA. All expression data are available at https://www.ncbi.nlm.nih.gov/geo/query/acc.cgi?acc=GSE173644.
